## Supplementary Data for "A single-cell gene regulatory network inference method for identifying complex regulatory dynamics across cell phenotypes"

### This PDF file includes:

Figs. S1 to S8

Tables S1 to S2

Commands S1 to S3

### Other Supplementary Materials for this manuscript include the following:

Excel file S1 to S2

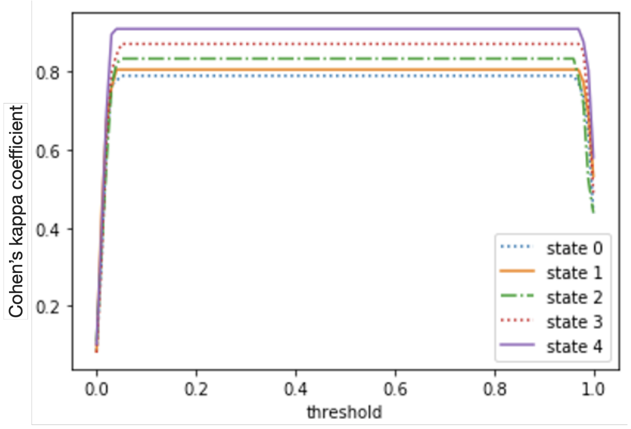


**(A)** SimiC

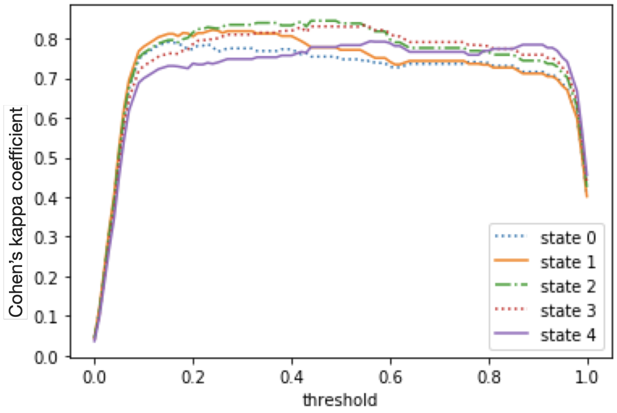


**(B)** c.LASSO

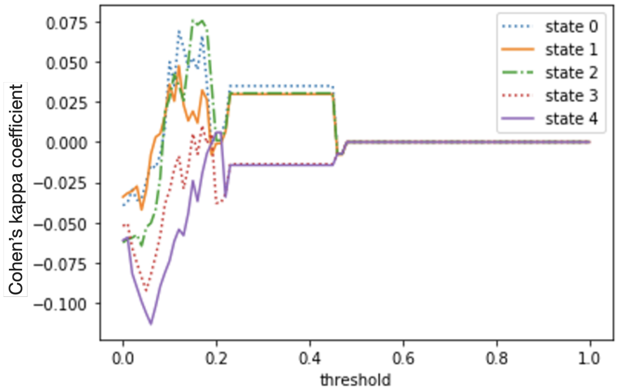


**(C)** SINCERITIES

**Fig. S1: Cohen's kappa coefficient for the 5 states of the generated synthetic dataset** obtained with **A:** SimiC, **B:** LASSO run in all states combined (denoted as c.LASSO) and **C:** SINCERITIES, for different thresholds. The threshold is used to convert the weight of the inferred edges into $0, +1$ and $-1$. Specifically, for a threshold $t$, all weights above $t$ are converted to $+1$, all weights below $-t$ to $-1$, and the rest to $0$. We observed that whereas the Cohen's kappa coefficient obtained with SimiC and c.LASSO is stable with the used threshold, that is not the case for SINCERITIES.

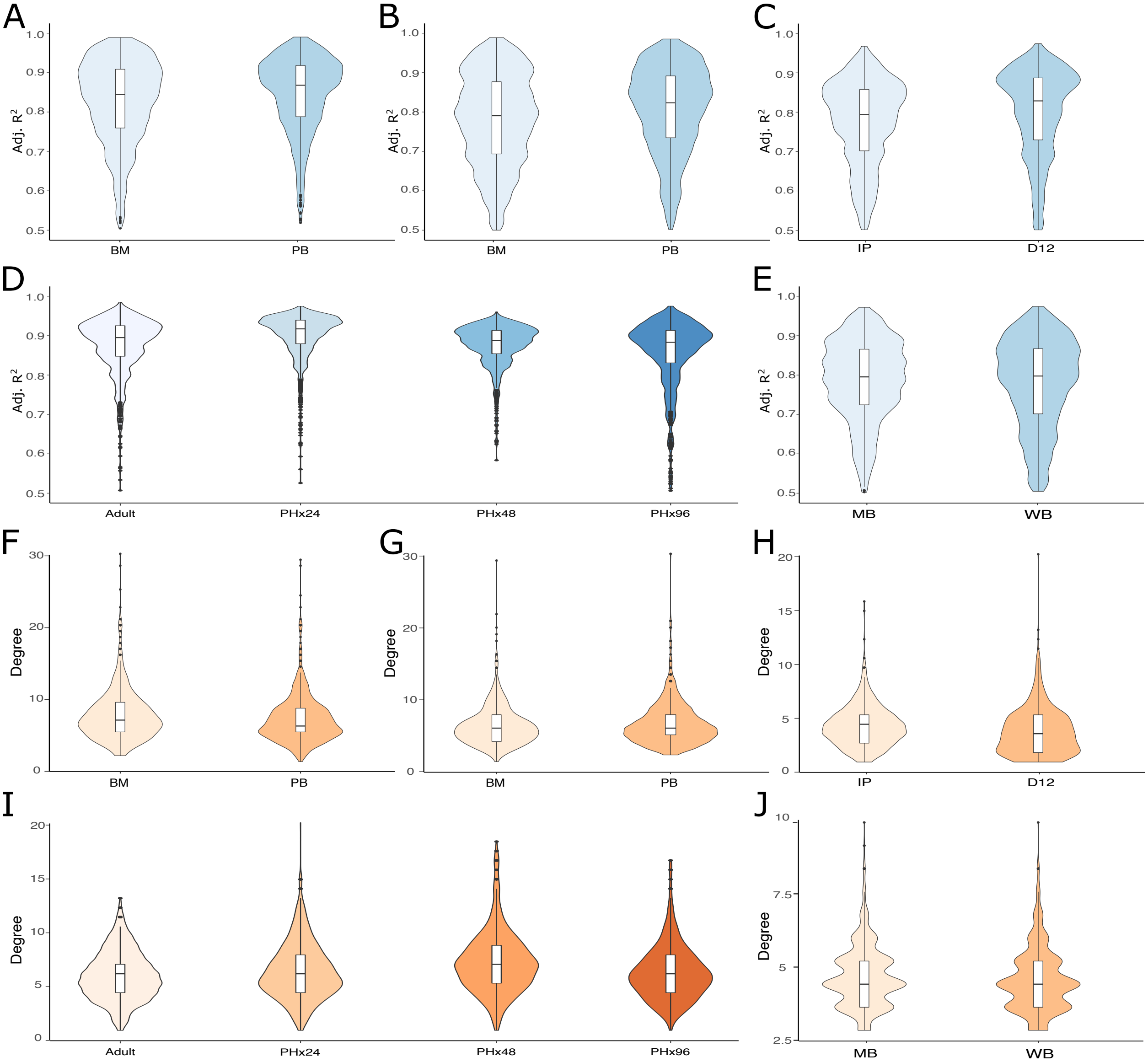


**Fig. S2: Goodness of fit (measured by the adjusted $R^2$) and number of transcription factors regulating the target genes for all the analyzed real datasets.** Violin plots (colored in blue) showing the distribution of the adjusted $R^2$ per target gene across the phenotypes on **A:** monocytes coming from either bone marrow (BM) or peripherial blood (PB) (*31*); **B:** CD4+ T-lymphocytes coming from either BM or PB (*31*); **C:** CAR T cells isolated from the infusion product (IP) as well as from PB at the expansion peak after treatment (D12) (*45*); **D:** hepatocytes sequenced at different timepoints (adult, PHx24, PHx48 and PHx96) after partial hepatectomy (*55*); **E:** bee brain cells coming from either WB or MB (*60*). The adjusted $R^2$ measures the goodness of fit between the true target genes' expressions and the ones computes with SimiC's inferred GRNs on the test data (which corresponds to about 20% of the data). Violin plots (colored in orange) showing the distribution of the number of transcription factors (TFs) regulating each target gene across the phenotypes on **F:** monocytes coming from either BM or PB (*31*); **G:** CD4+ T-lymphocytes coming from either BM or PB (*31*); **H:** CAR T cells at IP and D12 (*45*); **I:** hepatocytes sequenced at timepoints adult, PHx24, PHx48 and PHx96 (*55*); **J:** bee brain cells coming from either WB or MB (*60*). We observed that the median adjusted $R^2$ was above 0.8 in all cases, and the average number of transcription factors (TFs) regulating a given target gene was generally below 10.

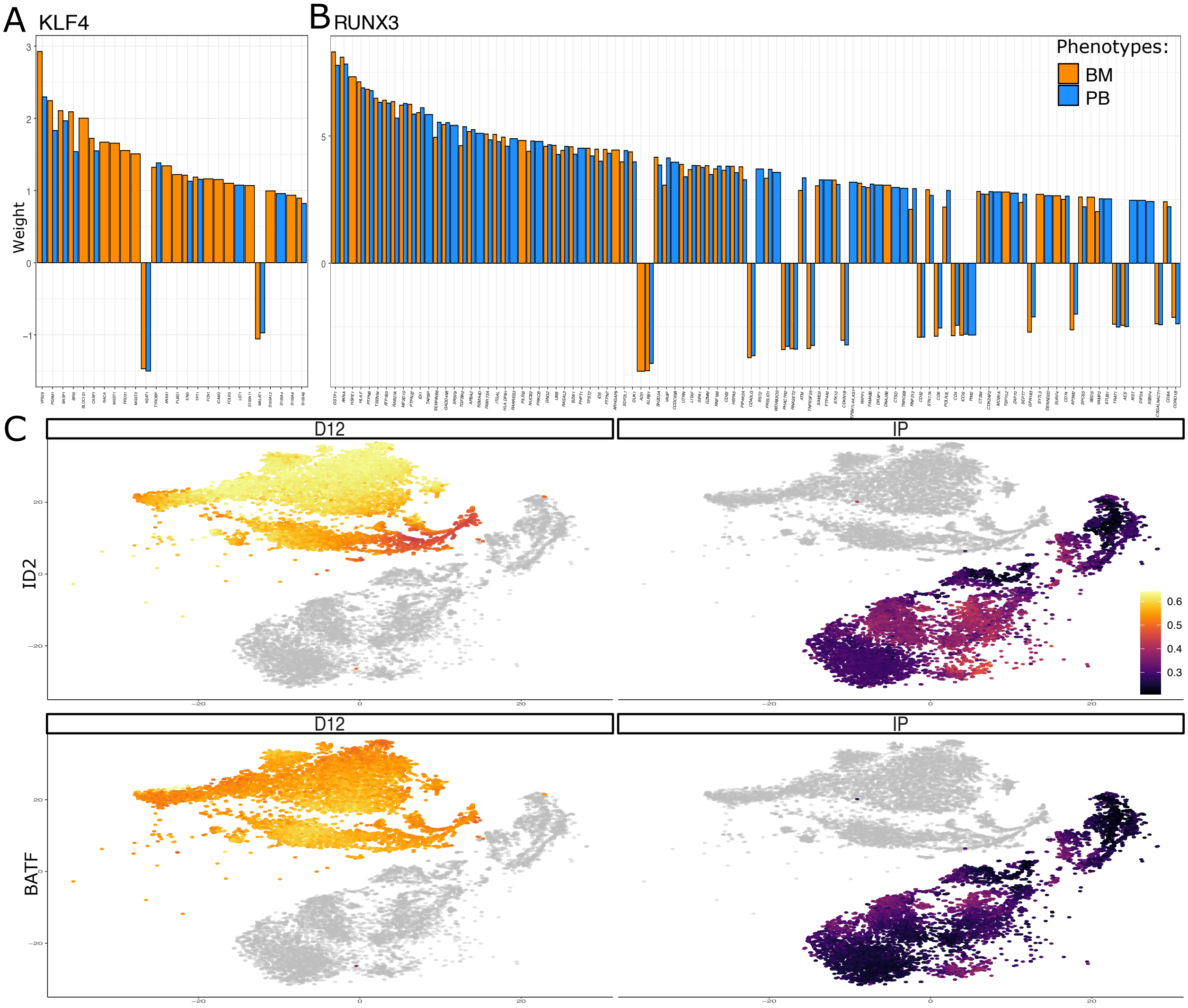


**Fig. S3: Additional plots regarding the assigned weights of the KLF4 and RUNX3 regulons from monocyte and CD4+ T-lymphocytes, respectively, as well as the activity of the regulons ID2 and BATF across the different CAR T cells.** Barplot showing the assigned weights to the target genes on the **A:** KLF4 regulon from monocytes coming from BM or PB (*31*) and **B:** on the RUNX3 regulon from the CD4+ T-lymphocytes coming also from either BM or PM (*31*). **C:** tSNE depicting the activity of the regulons ID2 and BATF across the different cells on the two different phenotypes (IP and D12) of the CAR T cells (*45*). These regulons show higher activity at D12 as compared to IP.

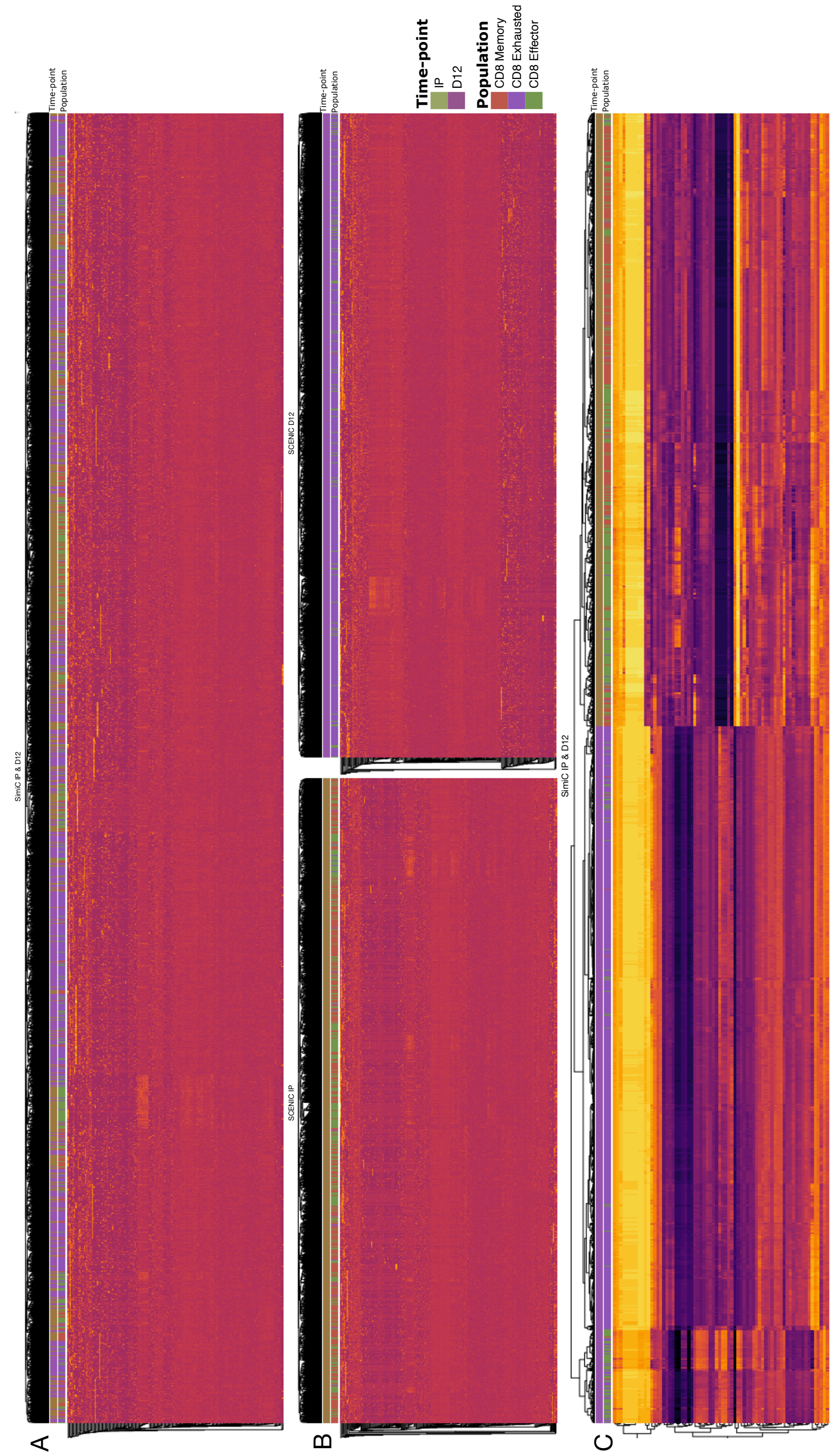


**Fig. S4: Performance comparison of SCENIC and SimiC on the CAR T dataset with phenotypes IP and D12 (*45*).** Similarly to SimiC, SCENIC computes a score per cell and transcription factor. We then looked at the results from clustering the cells based on their AUC score for SCENIC, and on their regulatory activity score for SimiC. For each cell, we specify the cell state: CD8 Memory, CD8 Exhausted, and CD8 Effector. **A:** Cell clusters using the AUCs calculated by SCENIC on the whole CAR T dataset (i.e., without differentiating between phenotypes IP and D12); **B:** Cell clusters using the AUCs calculated by SCENIC on each phenotype (IP and D12) separately; **C:** Cells clusters using the regulatory activity calculated by SimiC on the whole CAR T dataset. Note that SimiC jointly infers a GRN per phenotype (IP and D12). The heatmaps reveal that SCENIC's generated scores are not able to cluster cells based on their cell state (CD8 Memory, CD8 Exhausted, and CD8 Effector), whereas SimiC's scores can.

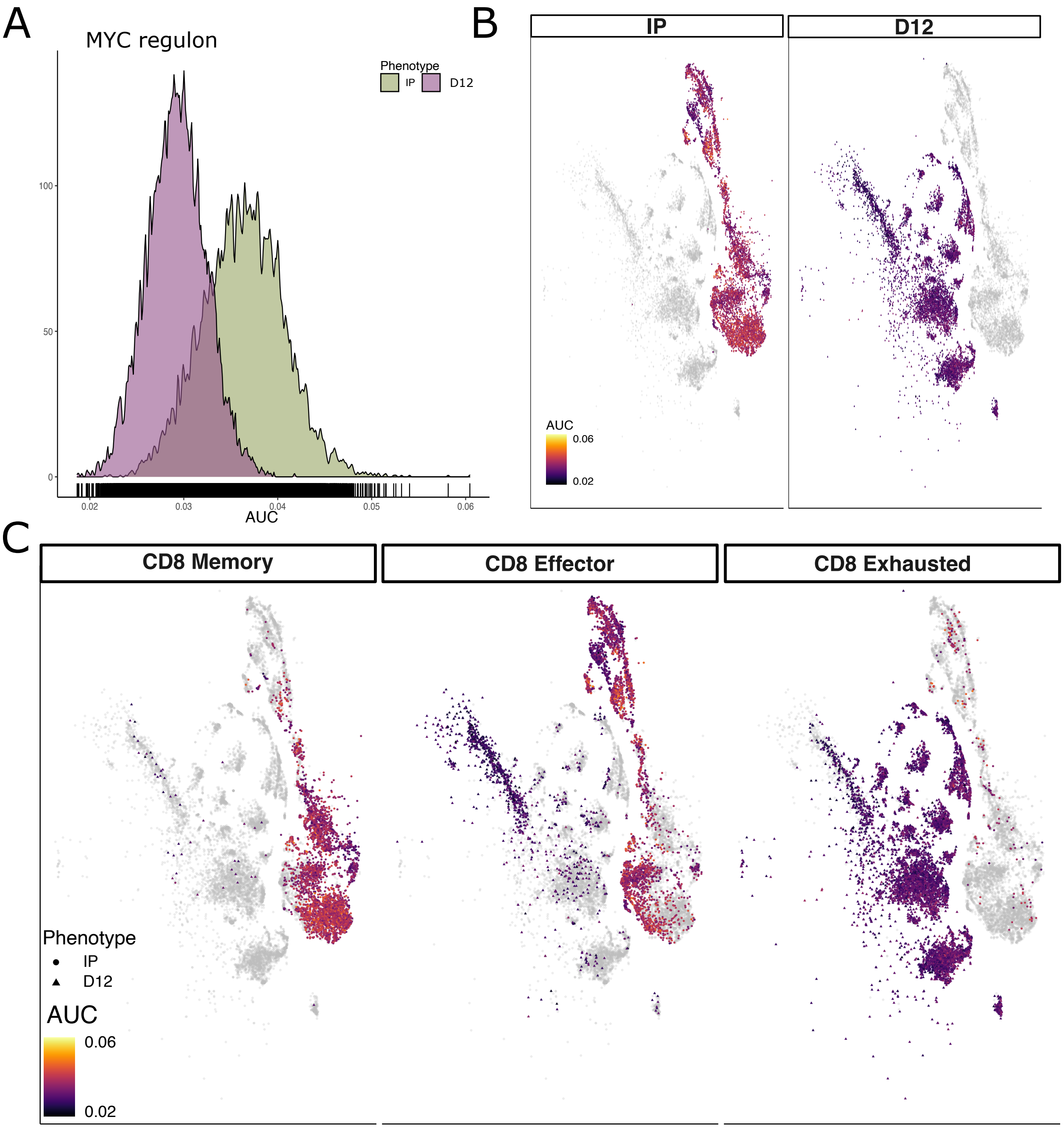


**Fig. S5: Analysis of the AUC scores computed by SCENIC for MYC regulon on the CAR T cell dataset (*45*).** The CAR T dataset contains two phenotypes (IP and D12), as well as three cell states (CD8 Memory, CD8 Exhausted, and CD8 Effector). **A:** Densities of the AUC score calculated by SCENIC for the two phenotypes IP and D12; **B:** tSNE visualization of the CAR T cells colored by the AUC score computed by SCENIC for the MYC regulon; **C:** tSNE visualization of the CAR T cells separated by cell state (CD8 Memory, CD8 Effector, and CD8 Exhausted), colored by the regulatory activity score computed by SCENIC for the MYC regulon. We show that while the AUC scores computed by SCENIC provide similar biological insights to that of SimiC's regulatory activity scores, contrary to SimiC, they were not able to recapitulate cell proliferation dynamics via the MYC regulon.

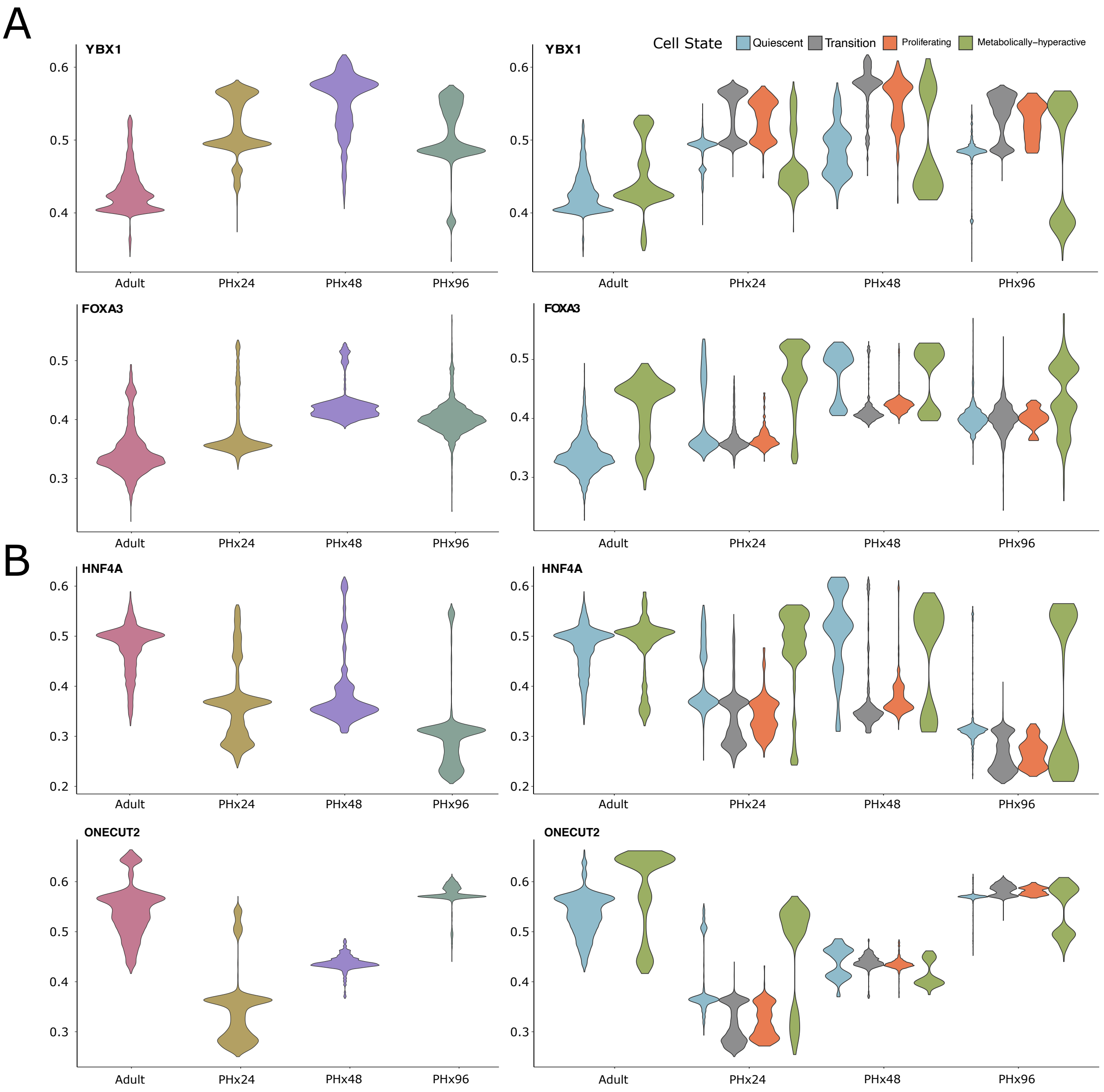


**Fig. S6: Additional results of SimiC for the hepatocyte dataset at timepoints adult, PHx24, PHx48 and PHx96 of a regenerating liver (*55*).** **A:** Violin plots showing the activity score distribution of the YBX1 and FOXA3 regulons for each timepoint, demonstrating their initiation-progression pattern. The activity score is also split by cell state; **B:** Violin plots showing the activity score distribution of the HNF4A and ONECUT2 regulons for each timepoint, demonstrating their termination-rematuration pattern. The activity score is also split by cell state.

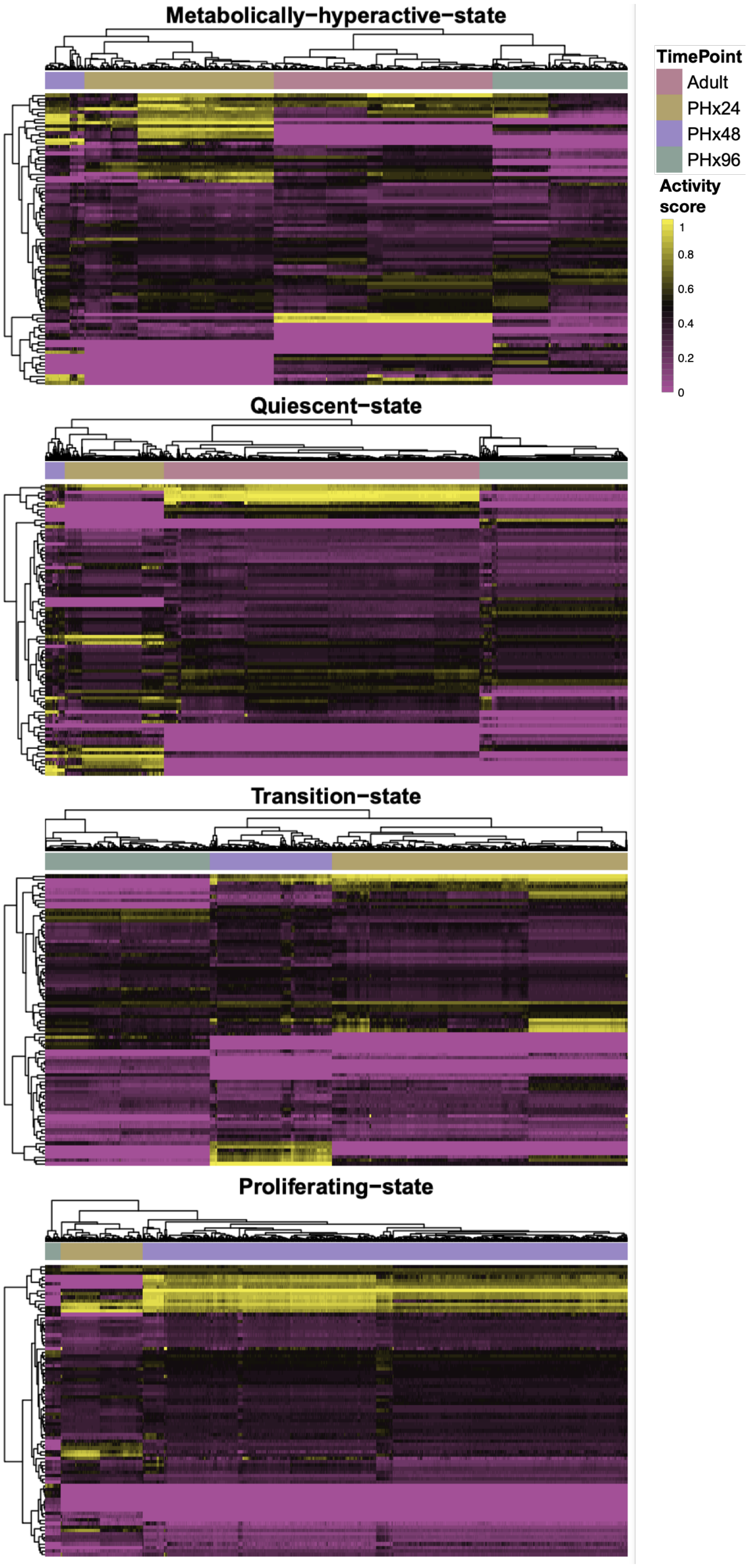


**Fig. S7: Clustering of regulon activity scores computed with SimiC for the hepatocyte dataset.** For hepatocytes sequenced at timepoints adult, PHx24, PHx48 and PHx96 of a regenerating liver (*55*), we show the heatmaps, including the clustering, of the regulon activity scores for each cell and for each TF, separated by cellular states. Note that cells cluster accurately by timepoint for all the different cell states, showing that SimiC's regulon activity score is able to capture timepoint-specific regulon activities for each cell state.

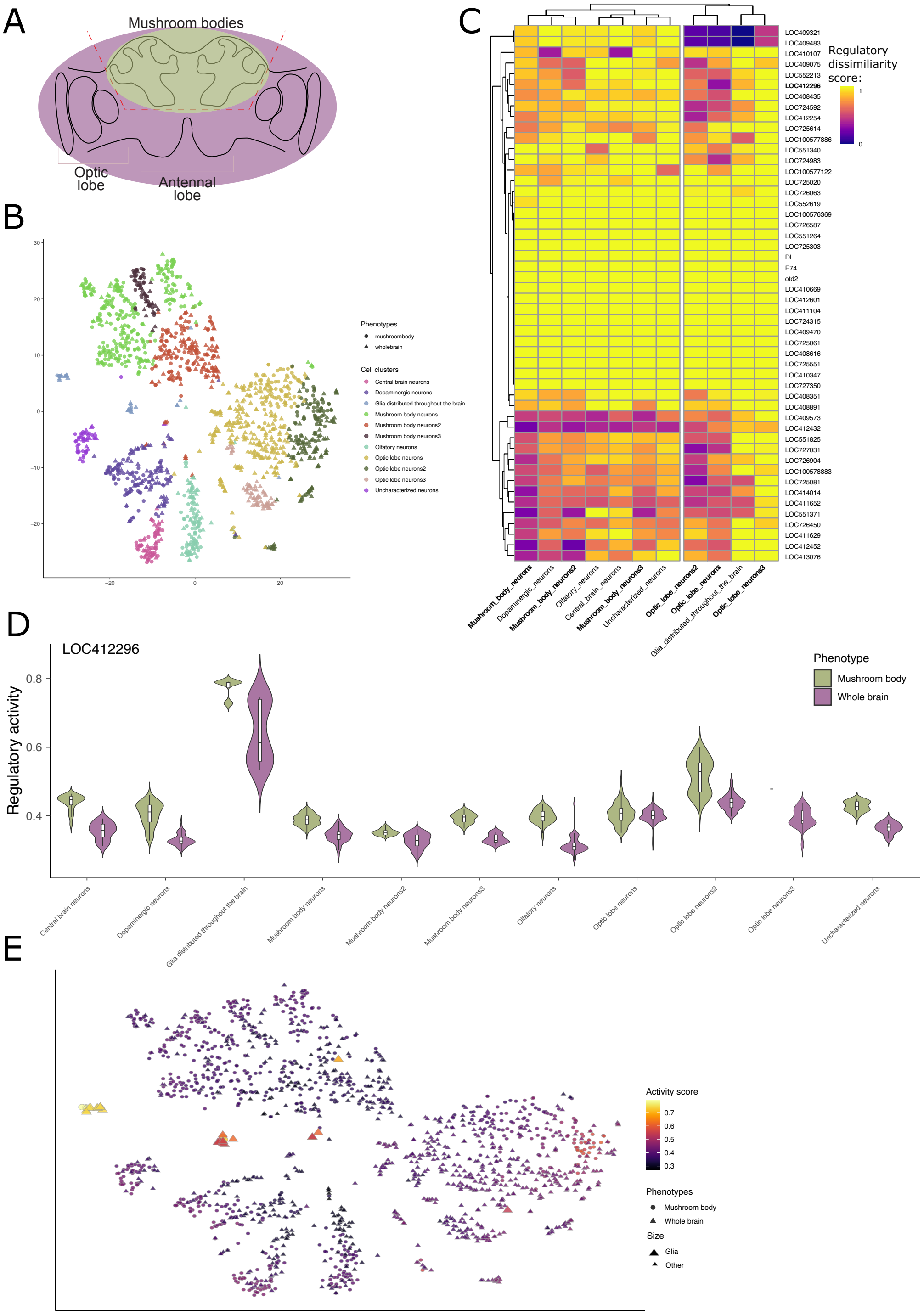

**Fig. S8: Additional results of SimiC on the bee brain dataset containing cells coming from either whole-brain (WB) or mushroom body (MB) (*60*).** **A**: Scheme depicting the base anatomy of the bee brain, and the line (dashed red) marking the surgical separation of the MB; **B:** tSNE visualization of the cells colored by cell state and shaped by the phenotype; **C:** Heatmap depicting the regulatory dissimilarity score across the whole brain and the mushroom body, for each cell state; **D:** Violin plot showing the regulatory activity of the regulon LOC412296 on the different cell clusters, on the mushroom body and whole brain. **E:** tSNE visualisation of the bee brain cells colored by the regulatory activity score of the regulon LOC412296. Shapes correspond to the two studied phenotypes and the size highligths the cells bellonging to the glia distributed throughout the brain.

| Transcription Factor | Odds ratio | Targets inferred by SimiC: With CHIP-Seq evidence | Without CHIP-Seq evidence | Transcription Factor | Odds ratio | Targets inferred by SimiC: With ChipSeq evidence | Without ChipSeq evidence |
|---|---|---|---|---|---|---|---|
| AHR | 2.4209334 | 85 | 18 | MEF2A | 18.4331055 | 4 | 1 |
| ATF3 | 7.4978883 | 26 | 2 | MEF2C | 5.9282092 | 37 | 11 |
| ATF4 | 2.9630598 | 15 | 19 | MXD4 | 1.1189897 | 3 | 5 |
| BACH1 | 4.3775359 | 49 | 25 | NFE2 | 5.4219379 | 164 | 30 |
| BCL6 | 4.5304360 | 114 | 25 | NFE2L2 | 0.9668978 | 4 | 6 |
| BHLHE40 | 0.9505501 | 4 | 5 | NFIL3 | 3.6513062 | 138 | 59 |
| CEBPA | 7.5685966 | 95 | 9 | NFKBIA | 1.2789404 | 3 | 261 |
| CEBPB | 3.8799309 | 154 | 17 | NFKBIZ | 3.3213372 | 144 | 87 |
| CEBPD | 6.1966466 | 213 | 61 | NONO | 9.2076705 | 57 | 15 |
| CUX1 | 3.5059642 | 107 | 80 | NR4A1 | 3.9580502 | 36 | 32 |
| ELF1 | 6.2987322 | 268 | 42 | POU2F2 | 4.6308920 | 319 | 52 |
| ETV6 | 0.9902268 | 2 | 8 | RBPJ | 2.1634393 | 36 | 18 |
| FLI1 | 4.3930636 | 90 | 15 | RUNX1 | Inf | 24 | 0 |
| FOS | 3.5783291 | 192 | 35 | RXRA | 3.4971783 | 17 | 6 |
| FOSL2 | 3.7329709 | 140 | 43 | SFPQ | 6.8906385 | 43 | 37 |
| GTF3A | 0.0000000 | 0 | 186 | SOX4 | 2.9117559 | 16 | 14 |
| HHEX | 0.0000000 | 0 | 29 | SPI1 | 5.2559537 | 437 | 32 |
| HIF1A | 5.3150490 | 99 | 12 | STAT1 | 3.2276457 | 62 | 19 |
| IKZF1 | 13.6800635 | 55 | 4 | STAT2 | 6.4613131 | 13 | 38 |
| IRF1 | 3.6048313 | 6 | 3 | STAT3 | 3.6470812 | 8 | 1 |
| JARID2 | 1.6326102 | 2 | 3 | TCF25 | 2.4640241 | 1 | 4 |
| JDP2 | 0.0000000 | 0 | 8 | TFDP1 | 3.7066129 | 102 | 101 |
| JUN | 4.6938915 | 37 | 3 | TRPS1 | 0.0000000 | 0 | 8 |
| JUNB | 3.4277315 | 348 | 131 | USF2 | 4.0261124 | 10 | 4 |
| JUND | 3.5951367 | 113 | 21 | XBP1 | 1.9454443 | 189 | 62 |
| KLF10 | 3.7823874 | 118 | 85 | YBX1 | 1.9573284 | 40 | 637 |
| KLF13 | 1.3175562 | 3 | 9 | YY1 | 8.6304982 | 64 | 4 |
| KLF3 | 8.4404662 | 4 | 33 | ZBTB7A | 3.2321853 | 26 | 10 |
| KLF4 | 2.9737157 | 19 | 9 | ZBTB7B | 0.0000000 | 0 | 1 |
| KLF6 | 5.3682047 | 60 | 17 | ZEB2 | 4.4777216 | 102 | 34 |
| LYL1 | 4.2432598 | 30 | 73 | ZNF467 | 2.7657130 | 78 | 80 |
| MAFB | 2.9669603 | 112 | 137 | ZNF524 | 2.6631704 | 3 | 5 |
| MAZ | 2.3720738 | 55 | 16 | | | | |

**Table S1: ChiP-Seq evidence for the monocyte dataset.** Results of the odds ratio test between each regulon and empirical TF-target binding evidence obtained from ChIP-Seq data, on the monocytes dataset (*31*).

| Transcription Factor | Odds ratio | Targets inferred by SimiC: With CHIP-Seq evidence | Without CHIP-Seq evidence | Transcription Factor | Odds ratio | Targets inferred by SimiC: With ChipSeq evidence | Without ChipSeq evidence |
|---|---|---|---|---|---|---|---|
| ARID4B | 2.4690861 | 11 | 7 | NCOR1 | 5.6515240 | 27 | 7 |
| ARID5B | 4.5884187 | 66 | 67 | NFKB1 | 0.6646428 | 2 | 5 |
| ATRX | 2.6881720 | 1 | 18 | NFKB2 | 0.0000000 | 0 | 1 |
| BATF | 5.5085389 | 65 | 27 | NFKBIA | 0.0000000 | 0 | 147 |
| BCL11B | 13.5380037 | 76 | 12 | NFKBIZ | 15.9520334 | 8 | 1 |
| BHLHE40 | Inf | 2 | 0 | NONO | 4.8289855 | 4 | 2 |
| CEBPB | Inf | 8 | 0 | NR3C1 | 2.6184494 | 16 | 3 |
| CREM | 8.2261561 | 6 | 1 | PBX2 | Inf | 1 | 0 |
| ELF1 | 8.4172264 | 43 | 5 | POU2F2 | Inf | 9 | 0 |
| ETS1 | 7.6077545 | 122 | 19 | PRDM1 | 5.0717411 | 8 | 1 |
| FLI1 | Inf | 2 | 0 | RUNX2 | 0.9345519 | 7 | 7 |
| FOS | 3.1939239 | 108 | 22 | RUNX3 | 4.8112994 | 103 | 39 |
| FOSL2 | 1.7113995 | 6 | 4 | SATB1 | 3.2277012 | 26 | 199 |
| FOXP1 | 6.8588888 | 94 | 13 | SFPQ | 3.1681276 | 38 | 71 |
| GATA3 | 5.9537226 | 69 | 8 | SMAD3 | Inf | 1 | 0 |
| GTF3A | 0.0000000 | 0 | 109 | SP140 | Inf | 1 | 0 |
| ID3 | 0.0000000 | 0 | 1 | STAT1 | 9.3820055 | 19 | 2 |
| IKZF1 | 4.9706751 | 40 | 8 | STAT3 | 11.8606038 | 26 | 1 |
| IRF1 | 5.3105098 | 123 | 42 | TCF25 | 2.3480087 | 5 | 21 |
| JUN | 2.9685132 | 39 | 5 | TCF7 | 3.9431971 | 170 | 55 |
| JUNB | 3.6435228 | 40 | 14 | THAP11 | 0.0000000 | 0 | 1 |
| JUND | 6.6351428 | 129 | 13 | TSC22D4 | 2.5602017 | 15 | 113 |
| KLF13 | 6.5932959 | 10 | 6 | UBTF | 3.3198623 | 4 | 3 |
| KLF3 | 3.3015387 | 1 | 21 | XBP1 | 2.0203395 | 73 | 23 |
| KLF6 | 4.6004665 | 252 | 84 | YBX1 | 1.8544620 | 31 | 519 |
| LEF1 | 3.6710235 | 90 | 165 | YY1 | 0.8075069 | 3 | 2 |
| MAF | 2.9986765 | 15 | 3 | ZBTB20 | Inf | 1 | 0 |
| MAX | Inf | 4 | 0 | ZBTB7A | 3.2324393 | 26 | 10 |
| MAZ | 4.8788522 | 211 | 30 | ZNF331 | 0.0000000 | 0 | 27 |
| MXD4 | 4.0401001 | 174 | 81 | ZNF394 | Inf | 1 | 0 |
| MXI1 | 3.1115985 | 27 | 13 | ZNF75A | 0.0000000 | 0 | 1 |
| MYC | 3.4385698 | 98 | 7 | | | | |

**Table S2: ChiP-Seq evidence for the CD4+ T-lymphocytes dataset.** Results of the odds ratio test between each regulon and empirical TF-target binding evidence obtained from ChIP-Seq data, on the CD4+ T-lymphocytes dataset (*31*).

### S1: Commands used to obtain the results for SCENIC

```
import scanpy as sc
import numpy as np
import loompy as lp

#Load the data
adata = sc.read_csv(cart_data_path, delimiter = '\t',
                    first_column_names=True)

# compute the number of genes per cell (computes 'n_genes' column)
sc.pp.filter_cells(adata, min_genes=0)

# mito and genes/counts cuts
mito_genes = adata.var_names.str.startswith('MT-')

# for each cell compute fraction of counts in mito genes vs. all genes
adata.obs['percent_mito'] = np.ravel(np.sum(
    adata[:, mito_genes].X, axis=1)) / np.ravel(np.sum(adata.X,
    axis=1))

# add the total counts per cell as observations-annotation to adata
adata.obs['n_counts'] = np.ravel(adata.X.sum(axis=1))

sc.pp.filter_cells(adata, min_genes=200)
sc.pp.filter_genes(adata, min_cells=3)
adata = adata[adata.obs['n_genes'] < 4000, :]
adata = adata[adata.obs['percent_mito'] < 0.15, :]

row_attrs = {
"Gene": np.array(adata.var_names),
}
col_attrs = {
"CellID": np.array(adata.obs_names),
"nGene": np.array(np.sum(adata.X.transpose()>0, axis=0)).flatten(),
"nUMI": np.array(np.sum(adata.X.transpose(),axis=0)).flatten(),
}

lp.create("cart_filtered.loom",adata.X.transpose(),row_attrs,
col_attrs)
# End of the Python commands, the rest should be executed on the terminal.

# generate the GRN
! pyscenic grn --num_workers 20 --output adj.tsv \
--method grnboost2 cart_filtered.loom  hs_hgnc_tfs.txt

# calculate the possible regulons
! pyscenic ctx adj.tsv  hg38__refseq-r80__10kb_up_and_down_tss.mc9nr.feather \
--annotations_fname motifs-v9-nr.hgnc-m0.001-o0.0.tbl \
--expression_mtx_fname cart_filtered.loom  --mode "dask_multiprocessing" \
--output reg.csv  --num_workers 20  --mask_dropouts

# calculate the cellular enrichment
! pyscenic aucell  PBMC10k_filtered.loom  reg.csv \
--output cart_SCENIC.loom  --num_workers 20
```

### S2: Commands used to obtain the results for SINCERITIES

```
####################
# PACKAGES required:
# glmnet
# ppcor
# cvTools
####################
library(glmnet)
library(ppcor)
library(cvTools)

# *** Data loading ***
uploading <- dget("SINCERITIES functions/uploading.R")
# DATA <- uploading('THP1 data/THP1_single_cell_data_EXCEL_no6_24_72_96.csv')
DATA <- uploading('synthetic_d50t20_5000_cells_w_header.csv')

# *** SINCERITIES ***

SINCERITIES_PLUS <- dget("SINCERITIES functions/SINCERITIES_PLUS.R")
result <- SINCERITIES_PLUS(DATA,noDIAG = 0,SIGN = 1,CV_nfolds = 10)
adj_matrix <- result$adj_matrix
SIGN <- 1

# Final ranked list
adj_matrix <- adj_matrix/max(adj_matrix)
final_ranked_predictions <- dget("SINCERITIES functions/final_ranked_predictions.R")
table <- final_ranked_predictions(adj_matrix,DATA$genes,SIGN=1,
directory = "Results", fileNAME='synthetic_d50t20_simic_cmp',saveFile = TRUE)
```

### S3: Commands used to obtain the results for SimiC

```
from simiclasso.clus_regression import simicLASSO_op
from simiclasso.weighted_AUC_mat import main_fn


p2df = 'ClonalKinetic.DF.pickle' # File containing the scRNA data
p2assignment = ClonalKinetics.clustAssign.txt' # File containing the cell phenotype, in this case, IP or D12
p2tf = 'ClonalKinetics.TFs.pickle' # File containing the list of the TFs to use for the regulon inference

similarity = True #Confirm the use of constrains
k_cluster = None #Do not use any clustering information from the scRNA dataset
num_TFs = -1    #Use all the TFs present on the TFs file
num_target_genes = -1 #Use all the targets to compute the regulons
max_rcd_iter = 100000
df_with_label = False

percent_of_target = 1 #Use all the targets to compute the regulatory activity
cross_val = True
p2saved_file_cross = 'ClonalKinetics_CrossVal_Ws.pickle'

# perform the cross-validation in order to select the optimal Lambda1 and Lambda2
simicLASSO_op(p2df, p2assignment, similarity, p2tf, p2saved_file_cross, \
              k_cluster, num_TFs, num_target_genes, max_rcd_iter = max_rcd_iter,\
              df_with_label = df_with_label, cross_val=cross_val)

#Once selected the optimal Lambda1 and Lambda2, fixe them and calculate the weights
cross_val = False
lambda1= 0.01
lambda2 = 0.01
p2saved_file = 'ClonalKinetics_L1'+str(lambda1)+'_L2'+str(lambda2)+'_Ws.pickle' #Weigths
p2AUC =        'ClonalKinetics_L1'+str(lambda1)+'_L2'+str(lambda2)+'_AUCs.pickle' #Auc

simicLASSO_op(p2df, p2assignment, similarity, p2tf, p2saved_file, \
              k_cluster, num_TFs, num_target_genes, max_rcd_iter = max_rcd_iter,\
              df_with_label = df_with_label, lambda1=lambda1, lambda2 = lambda2)

#Once the weights have been calculated, we can compute the regulatory activity for each regulon
main_fn(p2df, p2saved_file, p2AUC, percent_of_target = percent_of_target)
```
